# Contextualising UK moorland burning studies: geographical versus potential sponsorship-bias effects on research conclusions

**DOI:** 10.1101/731117

**Authors:** Lee E. Brown, Joseph Holden

**Affiliations:** water@leeds, School of Geography, University of Leeds, Leeds, LS2 9JT, UK

## Abstract

1. It has recently been claimed that geographical variability resulted in false conclusions from some studies examining the impacts of prescribed moorland burning, including the Effects of Moorland Burning on the Ecohydrology of River basins (EMBER) project. We provide multiple lines of evidence to contradict these claims and show that the EMBER results are reliable.
2. A systematic review of the literature also confirms that EMBER conclusions were not out of line with the majority of other published UK studies on responses to prescribed burning of *Sphagnum* growth/abundance, soil properties, hydrological change, or peat exposure and erosion.
3. We suggest that sponsorship-bias is associated with some recent research conclusions related to moorland burning. Thus, it is of grave concern when sponsorship or other potential conflicts of interest are not declared on publications related to moorland burning.
4. We show that sponsorship and other conflicts of interest were not declared on a recent publication that criticised the EMBER project, thereby entirely undermining that critical assessment.
5. *Policy implications:* The EMBER findings are robust. Our study suggests that publications on moorland burning that have been funded by pro-burning groups should be treated with extreme caution by the policy community. Publications that have been shown to have failed to declare conflicts of interest from the outset, when first submitted to a journal, should be disregarded by the policy community because peer reviewers and editors may have been unable to evaluate those pieces of work properly.

## 1. Introduction

Widespread intensification of land management in the UK uplands in recent decades to support the driven-grouse shooting industry (Douglas et al., 2015, Yallop et al., 2006) has led to a situation in which claims and counter-claims about the effects of this practice are now common practice. These claims stem from issues such as driven-grouse moors being linked to the unexplained deaths or disappearance of protected species such as the Hen Harrier (Murgatroyd et al., 2019), mountain hare population changes (Watson and Wilson, 2019, Hesford et al., 2019) and changes in vegetation and catchment processes due to vegetation burning (Yallop et al., 2010, Holden et al., 2012, Noble et al., 2019, McCarroll et al., 2016, Vane et al., 2013). Ashby & Heinemeyer (2019) (hereafter “A&H”) recently added to the debate with their critique of four of the Effects of Moorland Burning on the Ecohydrology of River basins (EMBER) project papers published to date. A&H suggested that the EMBER work was problematic, proposing that geographical variation had not been considered. This critique follows recent notable debates on aspects of UK moorland burning (Brown et al., 2016, Davies et al., 2016a, Davies et al., 2016b, Douglas et al., 2016, Evans et al., 2019, Heinemeyer et al., 2019) and addendums to research papers due to a lack of transparency from some authors regarding competing interests (Marrs et al., 2019a). To date, there has been no detailed wider analysis of the funding source or competing interests of scientists contributing to these debates to understand if this is a broader issue that should be taken into account.

This article first provides contradictory evidence to recent claims by A&H that geographical variability contributes to false conclusions from moorland burning studies. We present evidence to show that A&H’s critique is fundamentally problematic due to: (a) unexplained errors in their analysis including an incorrect portrayal of where geography (linked to site and plot specific analyses) was incorporated in previous analyses; (b) their errors being exacerbated by the avoidance of more recent papers published since 2016 using EMBER data, and (c) a selective focus on soil surface temperature when sensors buried at 5cm depth also revealed significant warming effects. Second, from a systematic review of the literature, we show that EMBER conclusions were not out of line with the majority of other published studies on responses to burning of *Sphagnum* growth/abundance, soil properties, hydrological change or peat exposure and erosion. Third, using this systematic review alongside available evidence on research funders, we show that sponsorship-bias (Lesser et al., 2007, Lexchin, 2003) may be associated with some recent research conclusions related to moorland burning. We also suggest that sponsorship interests and other conflicts of interest are of concern when they are undeclared on research outputs (Ashby and Heinemeyer, 2019, Marrs et al., 2019b) and that this undermines the research conclusions from those studies.

## 2. Methods

### 2.1 Re-examination of Ashby & Heinemeyer (2019) claims regarding geographical effects

To assess the claim that altitude was linked with precipitation across the EMBER study sites, data from Table 2 in A&H were assessed using linear regression. The analysis was also repeated with the altitude data reported from the catchment outlet in the original EMBER papers (e.g. Brown et al., 2013). We also tested for association between water temperature data (Supplementary Table 1) used in Brown et al. (2013) and altitude at the catchment outlet where the datalogger was situated. To assess the potential for effects of local geographical position on river invertebrate communities, catchment size and runoff parameters from Holden et al. (2015) were fitted to the NMDS (non-metric multidimensional scaling) solution using the envfit procedure described in Brown et al. (2013). Recent EMBER-related papers (Aspray et al., 2017; Brown et al., 2019) ignored by A&H show that fine particulate organic matter (FPOM) from peat erosion following burning has significant negative effects on river ecosystem invertebrate communities. FPOM densities reported by Brown et al. (2013) were therefore tested for association with catchment size and altitude at the catchment outlet. Additionally, FPOM associations with rainfall totals for the month of sampling were analysed by compiling rainfall data from the modelled gridded precipitation records as used by A&H (Supplementary Table 2).

**Table 1.** EMBER project-related ecosystem properties that were considered in the systematic review, and how each property was classified.

| **Ecosystem property** | **Classification** |
| --- | --- |
| *Sphagnum* growth/abundance response to prescribed burning | + = positive response to burning reported<br>- = negative response to burning reported<br>0 = no or mixed response to burning reported |
| Avoidance of mean and/or maximum soil temperature increase by prescribed burning | + = decreased temperatures reported<br>- = increased temperatures reported<br>0 = no temperature change |
| Reduced peat exposure and/or erosion by prescribed burning | + = reduced bare peat and/or erosion<br>- = enhanced bare peat and/or erosion<br>0 = no or mixed response to burning reported |
| Aquatic invertebrate diversity response to prescribed burning | + = positive response to burning reported<br>- = negative response to burning reported<br>0 = no or mixed response to burning reported |
| Prescribed burning alteration to peat physical and/or chemical properties (including pore water chemistry) | Yes = some change in response to burning reported<br>No = no change in response to burning reported |
| Prescribed burning alteration to stream chemistry | Yes = some change in response to burning reported<br>No = no change in response to burning reported |
| Prescribed burning alteration to peatland hydrological function | Yes = some change in response to burning reported<br>No = no change in response to burning reported |

**Table 2.** Correlation statistics for five flow variables (reported by Holden et al. (2015)) and the NMDS solution presented in Brown et al. (2013). The two variables that were correlated significantly with the NMDS solution were not associated with altitude or catchment size (Supplementary Figure 1).

| Flow variable | $R^2$ fit to NMDS | $p$-value fit to NMDS |
| --- | --- | --- |
| Time from start of rain to peak flow | 0.006 | 0.88 |
| Time from peak rain to peak flow | 0.13 | 0.04 |
| Rainfall before rise in river stage | 0.06 | 0.25 |
| Rainfall before steep rise in hydrograph | 0.002 | 0.96 |
| Recession time | 0.29 | 0.002 |

While there are no records of EMBER ‘surface’ thermistors being exposed periodically to direct sunlight and therefore being artificially warmed as claimed by A&H, we nevertheless tested the effect of this possibility to determine if it alters EMBER conclusions. Statistical models were developed by Brown et al. (2015) to predict daily maximum soil temperature in plots burned 15+ years prior to the study. These models were then applied to predict temperatures of plots burned 2, 4 and 7 years previously, with outliers (hereafter ‘disturbances’) from predicted temperatures enabling an assessment of the magnitude of burning effects. Using the maximum temperature datasets, the top 10% of temperature disturbances were discarded (Supplementary Table 3) and the analysis re-run following the same methods described in Brown et al. (2015).

### 2.2 Systematic review to contextualize EMBER findings

The aim of the review was to determine whether or not the EMBER project findings were significantly out of line with studies undertaken before the project started as well as in recent years. We undertook a meta-analysis using a combination of Web of Knowledge and Google Scholar searches conducted between 28/06/19 and 02/07/19 to compile UK upland burning literature. These searches were supplemented with examination of reference lists from recent publications, including other systematic reviews (e.g. Glaves et al., 2013), and our own knowledge related to grouse moor management and prescribed vegetation burning research outputs. We initially rejected studies which were not based in the UK uplands, those focusing solely on wild, uncontrolled fire effects, review/opinion/comment papers or grey literature, and those with no obvious focus on environmental responses to grouse moor management. The initial searches and shortlisting produced a list of 112 potentially relevant, publically-accessible peer-reviewed papers (Supplementary Table 4).

From the shortlisted papers, both authors subsequently reviewed each paper together, focusing particularly on abstract, results and conclusion sections to categorise papers in terms of their relevance to seven ecosystem properties that were studied in the EMBER project (Table 1). For four of the properties, we considered that it was possible to classify responses to vegetation burning as positive, negative, or having no/mixed effects. We classified such responses when the authors of those papers either made clear statements suggesting there was a burning effect, and/or when they reported a statistically significant effect. For the other three ecosystem properties (soil physical/chemical properties, stream water chemistry, hydrology) we classified responses in terms of whether there was a change/difference (yes) or no change/difference (no) reported. This avoided a subjective value judgement, because many of these studies reported data without any clear statement as to whether the changes or differences could be deemed positive or negative for peatland function. Finer-scale properties for specific variables (e.g. pH, DOC, EC as part of the stream water chemistry group) were explored initially but returned low numbers of studies, hence our use of broader groupings. Overall, 61 of the 112 shortlisted papers were considered to be directly relevant for comparison with the EMBER project findings for at least one of the seven ecosystem properties (Supplementary Table 5). All retained papers that were found to be relevant to the first four ecosystem properties were also classified in terms of a net effect: + (only positive outcomes reported across the four properties), - (only negative outcomes reported), or 0 (either only 0 outcomes or a mixture of + and – outcomes reported).

### 2.3 Potential sponsorship bias

For each retained paper that was classified into the seven properties in Table 1, the acknowledgements or funding declaration section (where present) was used to determine which organizations had been declared as funders of the research, along with any relevant competing interests declared by the paper author(s). We then combined the papers into groups for further analysis: (1) those declaring funding links to groups which are considered to be supportive of prescribed burning as part of grouse moor management (Game & Wildlife Conservation Trust (GWCT), formerly the Game Conservancy Trust (GCT); The Heather Trust, The Moorland Association, plus landowners or estates involved in managing grouse shoots), and (2) other groups, where a position on grouse shooting is not available (e.g. government departments/agencies, research councils, universities, upland restoration groups such as Moor for the Future/Yorkshire Peat Partnership, water companies) and studies where no funding information was provided. We also examined datasets to determine if there was an association between funding and research conclusions for studies with: (3) government agencies that form national-scale environmental policy (e.g. Department of Agriculture for Northern Ireland, Department for Environment, Food and Rural Affairs (Defra) – formerly the Ministry of Agriculture, Fisheries and Food (MAFF), Natural England - formerly English Nature, Scottish Government, Scottish Natural Heritage) versus (4) no government agency acknowledgements.

Due to the relatively small number of observations available across ecosystem properties, Fisher’s Exact Test for count data was used to test associations between funding groups and research conclusions for each of the seven ecosystem properties and the net-effect using R v.3.5.2 (Supplementary Table 6). Pairwise comparisons were conducted using the Fisher multi-comparison test in the RVAideMemoire package (Hervé, 2019), and applying a Bonferroni correction. The test was unconditioned because rows and column totals varied. The test assumes independence of observations between each published study; this might not be the case for some long-term studies such as those reporting vegetation changes over time at the Moor House experimental plots in northern England (Milligan et al., 2018, Marrs et al., 2019b, Lee et al., 2013) but here we assumed them to be independent because each successive study reported new observations.

## 3. Results and discussion

### 3.1 Re-examination of Ashby & Heinemeyer (2019) claims regarding geographical effects

A&H suggested that the EMBER project results are unreliable because they were based on a correlative space-for-time approach with treatments located in geographically-separate and environmentally-distinct sites, and they further implied that this was not accounted for in the analysis of the data. This critique is odd because: (1) the analyses did examine numerous site-specific variables and differences; (2) the EMBER project included experimental Before-After-Control-Impact (BACI) manipulations of some key parameters so it was not solely a space-for-time project (e.g. Aspray et al., 2017, Brown et al., 2019); (3) A&H themselves have advocated the use of geographically separate study sites to defend their own research on moorland burning because “sampling across a wider area with climatic differences should be seen as an advantage, as it offers real and meaningful replication rather than providing detailed records for only one site” (Heinemeyer et al., 2019, page 2).

A&H suggested that slope was not accounted for in the EMBER papers. However, A&H made a fundamental mistake in their assessment of how EMBER studies incorporated slope. Four papers (Brown et al., 2013, Brown et al., 2015, Holden et al., 2015, Holden et al., 2014) stated that plot locations were determined based on topographic index (TI) categories so that across all ten catchments there were three groups of plot locations defined by the TI. The TI (Beven and Kirkby, 1979) is given as ln (*a* / tan β) where is *a* is slope length per unit contour width and β is the local topographic gradient. This is a much more logical approach for comparing treatment effects on soil hydrological function and associated soil properties than just using slope angle alone. A peatland plot location with the same slope angle would be expected to have a very different distribution of water-table fluctuations and saturation extent depending on the slope length draining to that point, as the upslope water supply for that plot would be greater for longer slopes (Holden, 2005). The TI is therefore used worldwide in hydrological and ecological studies to support environmental analysis and sample point selection as it incorporates both slope and slope length (e.g. Zinko et al., 2005, Li et al., 2018c, Mohamedou et al., 2019). Unfortunately, many blanket peatland management impact studies (e.g. Heinemeyer et al., 2018, Lee et al., 2013, Clay et al., 2009, Jonczyk et al., 2009) have neglected to recognise or factor in *a* and upslope contributing flow concepts to their study designs which makes interpretation of their findings difficult. A&H used an Ordnance Survey digital elevation model with 50 m × 50 m grid cells to estimate the slope of each of the EMBER plots and suggested that there was a significant difference in slope between unburnt (steeper) and burnt plots. However, their analysis suggests that the difference was <1°. First, given that EMBER plots were approximately 20 × 20 m, 1° must be well within the margin for error when using a 50 × 50 m digital elevation model across the UK uplands. Second, A&H should have established whether the effect size is meaningful (unlikely as blanket peatland is often found covering slopes up to 20° (Lindsay et al., 1988) and in extreme cases up to 30° (Ingram, 1967)). Third, even if an effect size of <1° was meaningful, in theory steeper plots would have a greater likelihood of deeper mean water-table depths with more variability than less steep plots. However, Holden et al. (2015) reported burnt plots had significantly deeper mean water-table depths and greater water-table variability than unburnt ones. This is the opposite of what A&H’s slope analysis would suggest. A similar point can be made about the potential effects of slope that A&H hint at (although they do not explain what effects there could be) for the macropore flow and hydraulic conductivity study by Holden et al (2014). Holden (2009) already established that more gentle peat slopes are associated with higher macropore flow and saturated hydraulic conductivity. Yet Holden et al. (2014) showed that plots with recent fire (prescribed or wildfire) had lowest macropore flow and saturated hydraulic conductivity, no matter whether they were on less steep, equal or steeper slopes than other treatments. Hence, this provides additional evidence that the managed burning effects were large and impacted the hydrological functioning of the peatland plots studied in the EMBER project, thereby strengthening the EMBER project’s conclusions.

A&H hinted that plots (by burn age or burnt/unburnt) were distributed variably with respect to aspect but they did not find any statistically significant effect. A&H noted that “elevation exerts a strong influence on precipitation which, in turn, affects peatland water tables and overland flow”. By inference they go on to suggest that our hydrological data from the ten EMBER case study sites are therefore problematic as the ten catchments were (unavoidably) in different locations. However, they show in their own analysis (Figure 2c in A&H) that there is no significant difference in elevation between the burnt and unburnt catchments (they report *p*=0.465). Furthermore, using the precipitation and elevation data presented by A&H in their Table 2, we find two major problems: (1) across the ten sites there was no significant relationship between elevation and mean monthly precipitation (using A&H’s mean elevation, *R*^2^ = 0.19, *p*=0.21, or using the catchment outlet elevation reported in our papers *R*^2^ = 0.11 *p* = 0.36); (2) A&H presented the same precipitation values for two catchments in both burned and unburned categories, with no explanation of the modelling errors that underpin this issue and thus their analysis overall. By including these non-independent gridded rainfall estimates twice in their analysis, A&H have arguably produced a misleading statistical output. With only n=4 for each land use, and using ANOVA as per A&H, the summary statistics for Figure 2b in A&H’s paper become *p*=0.07, *R*^*2*^ = 0.45.

**Figure 1.**
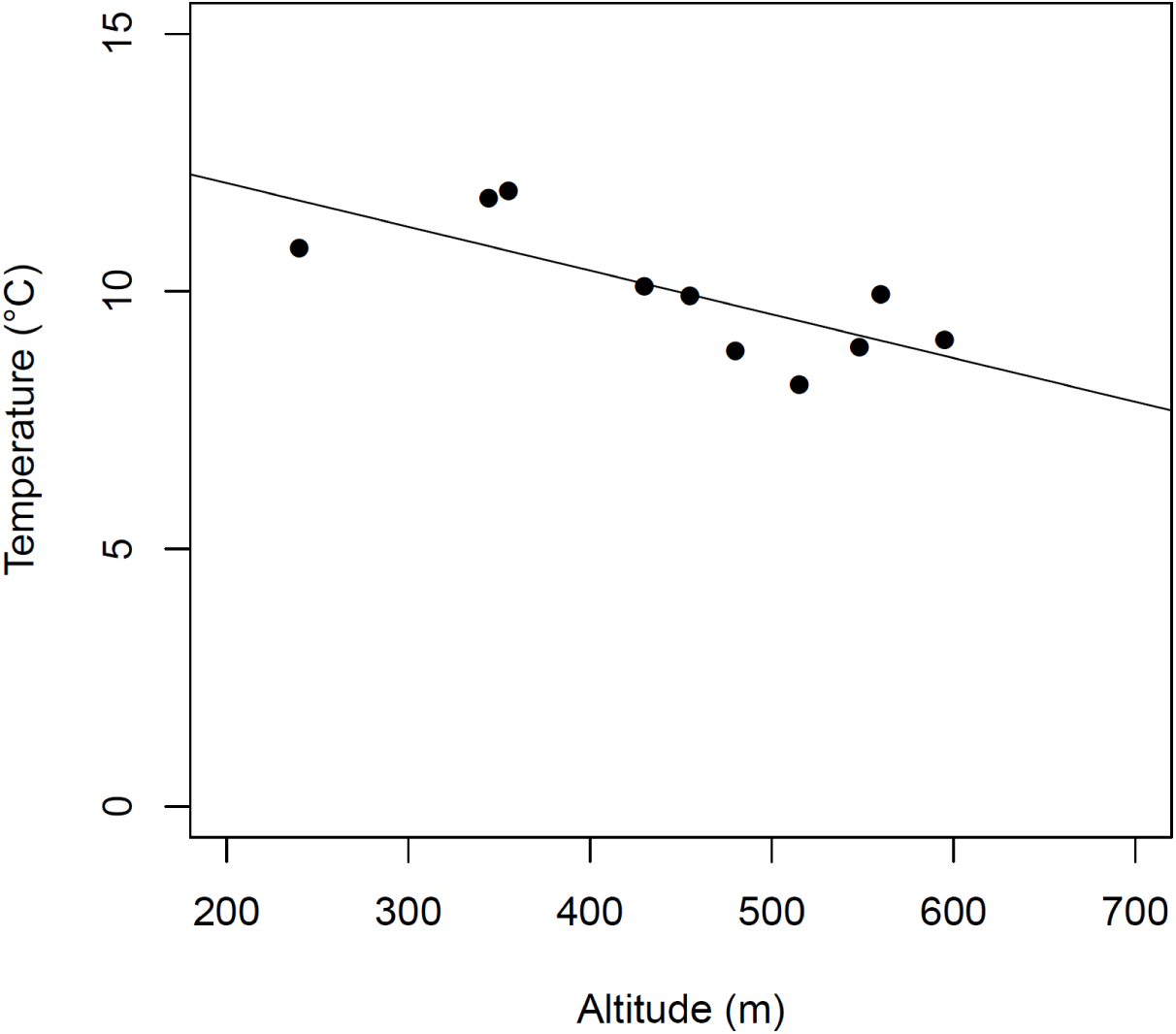
Relationship between altitude and mean river water temperature at the ten sites studied in Brown et al. (2013). [*R*^2^ = 0.56, *p*=0.013]. See Supplementary Table 1 for temperature data; altitude data were reported in the original paper.

**Figure 2.**
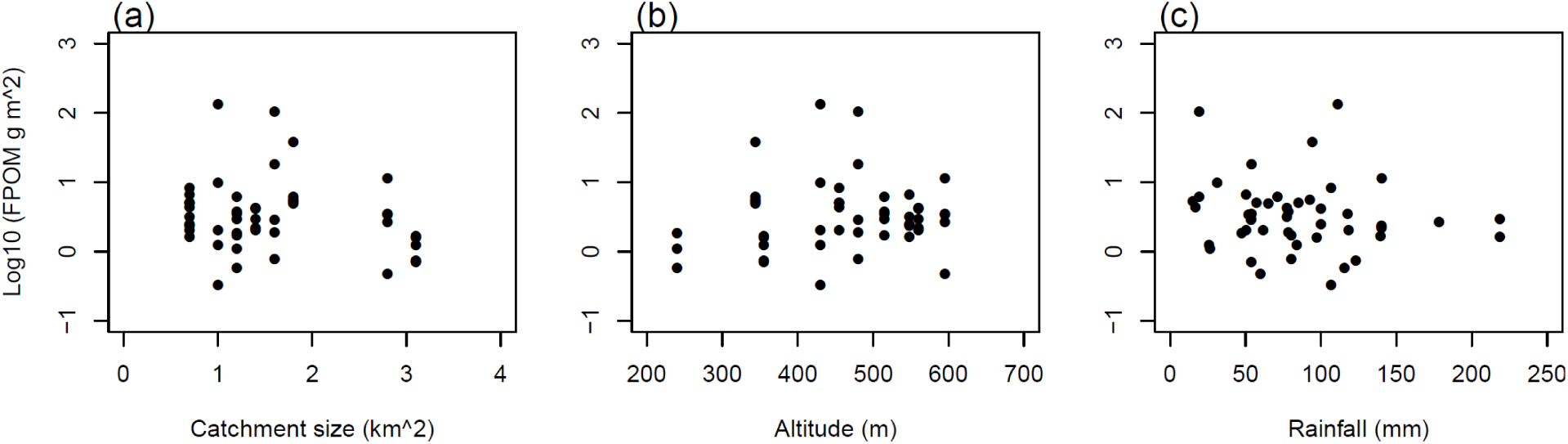
No relationships were evident between fine particulate organic matter (FPOM) densities and (a) catchment size (*R*^*2*^=0.007, *p*=0.57), (b) altitude (*R*^*2*^=0.001, *p*=0.81) or (c) precipitation during month of sampling (data sources as in A&H) (*R*^*2*^=0.005, *p*=0.63) in the EMBER rivers studied by Brown et al. (2013). See Supplementary Table 2 for FPOM and rainfall data; catchment size and altitude data were reported in Brown et al. (2013). Log transformation is used only for clarity of presentation; statistics presented are for untransformed data. For c, data sources from A&H were used for rainfall estimates and so it should be noted that for some sites the values were the same due to the gridded modelling errors associated with the analysis by A&H.

Contrary to unsubstantiated claims by A&H, Holden et al. (2015) clearly accounted for possible between-catchment rainfall effects in their analysis of storm event responses and provided a site by site breakdown of the results (e.g. Tables 3 and 4 within Holden et al., 2015). For example, *for every catchment* 25 different rainfall events were randomly selected from across the full range of rainfall event sizes experienced in that catchment, avoiding periods of snow and snowmelt (i.e. 250 rainfall events in total). Furthermore, contrary to the claims in their paper that suggested that EMBER did not control for site effects, A&H do actually point out that a statistical analysis of the EMBER data included site as a random factor within models when examining the effect of several environmental variables on vegetation (Noble et al., 2018). Curiously, A&H decided for themselves that this “*was not associated with the main EMBER project*”, despite the fact that the funding and acknowledgements of the Noble et al. (2018) paper are clearly attributed to the EMBER grant, three of the authors were from the EMBER team, and that the paper investigated the EMBER plots. Importantly, Noble et al. (2018 p565) already showed that “*the vegetation of the unburned sites shows a clear divide between North and South Pennine sites in the EMBER NMDS ordination in line with the two mire NVC [National Vegetation Classification – (Rodwell, 1991)] types they supported. However, this north–south divide is not apparent in the burned plots, and three of the five burned sites supported heath vegetation types associated with burning, grazing and atmospheric pollution (Elkington et al., 2001). This suggests that geographically variable vegetation community characteristics can be overridden by the effects of burning.”*

**Table 3.** Mean ±1 St. Dev *û*_t_ estimates for odd day maximum daily temperature predictions at the soil surface, with significance results from K-S tests for each EMBER age plot (B7+, B4 and B2 = plots burned more than 7, 4 or 2 years prior to measurement) relative to B15+ (plots last burnt more than 15 years prior to measurement) [^x^ = p>0.05; * = p<0.05; **p<0.01; ***p<0.001]. Values in square parentheses are Cliff’s δ estimates of effect size. See Brown et al. (2015) for original data analysis.

| Slope-position | B7+ | B4 | B2 |
| --- | --- | --- | --- |
| ***Topslope*** |  |  |  |
| Brown et al. (2015) | 0.63±2.37*<br>[0.15] | 1.88±3.65***<br>[0.35] | 2.77±4.49***<br>[0.42] |
| Top 10% $\hat{u}_t$ estimates removed | 0.09±1.65*<br>[0.17] | 1.09±2.72***<br>[0.38] | 1.78±3.36***<br>[0.45] |
| ***Midslope*** |  |  |  |
| Brown et al. (2015) | -0.21±1.99 | 1.46±3.50***<br>[0.26] | 1.04±2.44***<br>[0.26] |
| Top 10% $\hat{u}_t$ estimates removed | -0.57±1.74 | 0.65±2.53***<br>[0.28] | 0.55±1.95***<br>[0.29] |
| ***Footslope*** |  |  |  |
| Brown et al. (2015) | 0.89±3.46*<br>[0.12] | 0.30±2.65 | 2.27±4.66***<br>[0.31] |
| Top 10% $\hat{u}_t$ estimates removed | 0.09±2.42*<br>[0.11] | -0.31±1.81 | 1.17±3.13***<br>[0.32] |

**Table 4.**
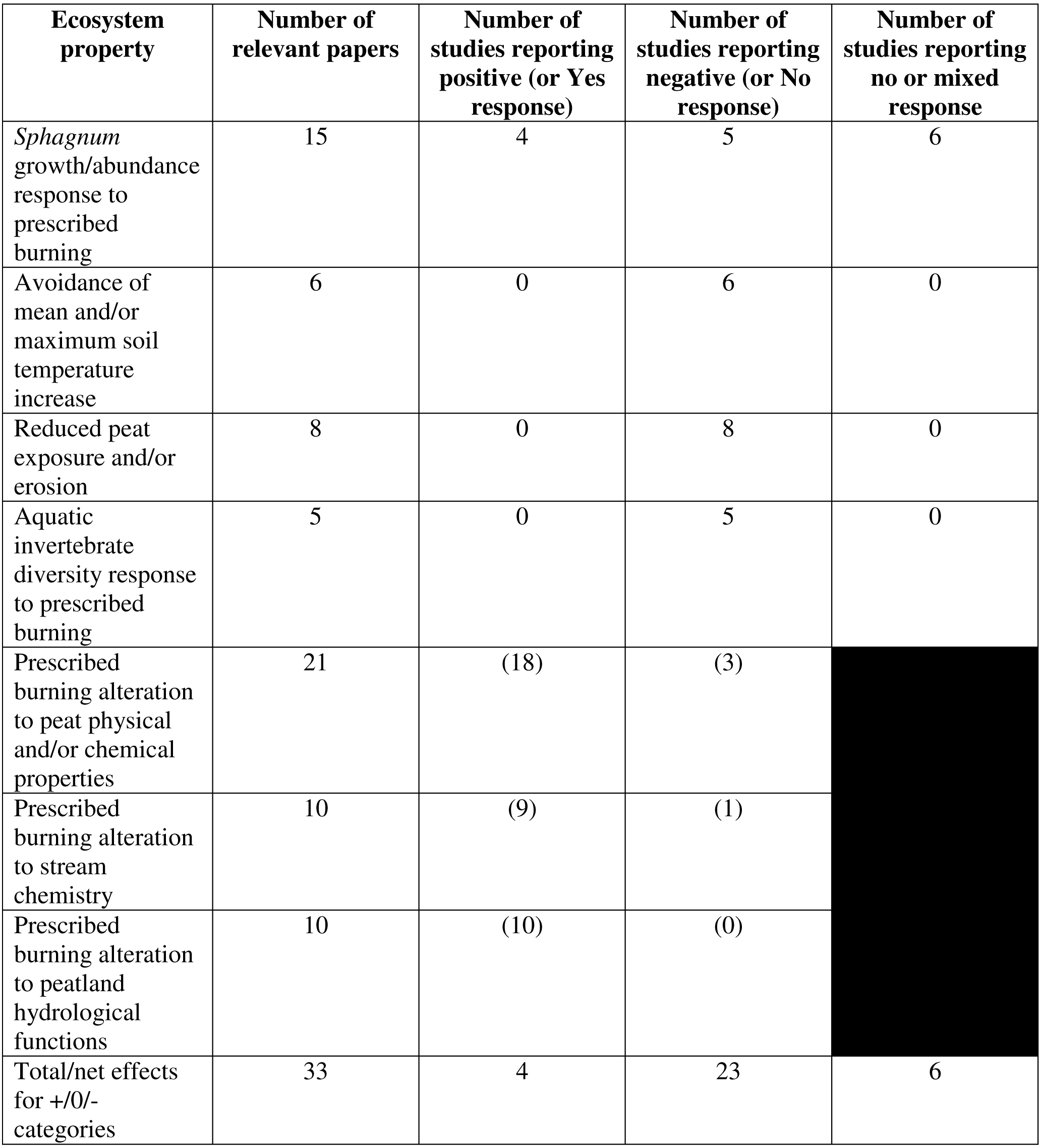
Summary statistics from the systematic literature review for seven ecosystem properties relevant to EMBER project papers.

A&H failed to acknowledge how site-based variables were considered in the analysis of data in other EMBER papers. In Brown et al. (2015), two study sites were used and it was explained clearly in the methods text that strong linear relationships were demonstrated between soil temperatures measured at burned site plots and those measured at the unburned site plots. Thus, any between-site differences in environmental parameters such as altitude or precipitation were implicit within the statistical relationships between the two sites. The strong linear relationships between the two sites were then used as the basis of examining effects *within* the burned site only. For Brown et al. (2013), A&H criticised the analysis for not considering between-site differences when analysing macroinvertebrate community and river habitat parameter responses to burning. Their suggestion is misleading because the analysis did include river water temperature data, which is associated strongly with altitude (Figure 1). Analyses already detailed in Brown et al. (2013) showed that water temperature was not associated significantly with the NMDS solution, nor was catchment size (*R*^2^ = 0.04; *p*=0.365). Rainfall was not incorporated into this analysis because rainfall-runoff relationships are modified by catchment processes, and river invertebrates would thus respond to flow changes rather than rainfall. Incorporation of mean flow metrics provided by Holden et al. (2015) suggest invertebrate communities may be associated with flow variability in our study, but where this is the case (Table 2) these flow-related changes are those associated strongly with vegetation burning and not with geographical variables (Supplementary Figure 1) as outlined by Holden et al. (2015).

Surprisingly, A&H suggested that the EMBER project was only a space-for-time study, ignoring two recent papers in which peat inputs were manipulated experimentally in both a real stream and a replicated streamside mesocosm facility, and incorporating a BACI design (Aspray et al., 2017; Brown et al., 2019). Vegetation removal as a result of fire often exposes the peat surface enhancing erosion by wind and precipitation, something which has been reported long before the EMBER project (Kinako and Gimmingham, 1980) and which is recognised explicitly as a problem associated with vegetation burning in the Heather & Grass Burning Code (Defra, 2007) and the Scottish Muirburn Code (Scottish Natural Heritage, 2017). These BACI experiments were designed to increase understanding of how peat particles that subsequently reach the river bed might explain the ecological differences that we presented in our earlier paper (Brown et al., 2013). Brown et al. (2013) demonstrated that fine particulate organic matter (FPOM) was significantly higher in burned river systems within the EMBER project and here we show that FPOM displayed no statistical relationships with either catchment size, altitude or precipitation (Figure 2). Both experiments (Aspray et al., 2017; Brown et al., 2019) implicated the build-up of peat particles as a major cause for change in the biodiversity and functioning of upland rivers.

A&H claimed that some of the soil-surface temperatures measured in the EMBER recently-burned plots (B2 - within two years of burning) could be due to measurement error caused by the particular type of sensor being located at the peat surface with parts exposed to direct sunlight. While Brown et al. (2015) used the term ‘surface’ for sensors placed shallowest in the soil profile, they also explained that the probes were placed horizontally within the top 1 cm of the peat-litter layer (i.e. not directly on the peat surface) and checked for position every three weeks. The study showed that maximum temperatures recorded at the most recent EMBER burn plots were similar to those reported by Kettridge et al. (2012) for Canadian peatlands in the aftermath of fire but at a similar latitude to our study sites, even though they used a different type of sensor (see discussion in Brown et al. 2015). Thus we have confidence in the temperature data from our study sites. If the EMBER temperature sensors were exposed to the sun periodically as claimed by A&H this provides no explanation for why: (i) significantly higher temperatures were recorded over the study period with sensors that were buried at 5cm depth in the most recently burned patches (B2 plots, burned less than two years prior to measurements) (Brown et al, 2015) - the surface must have been warmer also; (ii) exposure to sunlight cannot explain why the very lowest temperatures were also recorded in the B2 plots, which in turn would enhance soil ice formation and erosion processes (Li et al., 2018a, Li et al., 2018b); (iii) re-analysis of the peat temperature data from the EMBER plots with the top 10% of disturbance values removed match those presented in Brown et al. (2015), although the magnitude of the disturbances was, of course, reduced (Table 3). Notably though, removal of the top 10% of the highest temperatures increased the effect size for 6/7 burned plots where there was a statistically significant increase in temperature compared to B15+ plots (plots last burned more than 15 years prior to measurement). Thus, it can be concluded that even after exclusion of the most extreme (top 10%) temperature measurements from the EMBER dataset, vegetation removal with fire still increases maximum soil temperatures in the years that follow. In addition, recent experimental studies in other peatlands following vegetation burning have independently shown significant temperature increases measured at 2 cm depth (Grau-Andrés et al., 2019). Thus, there is a growing body of evidence that vegetation removal with fire modifies peatland soil thermal regimes in the years after fire.

### 3.2 Systematic review to contextualise EMBER findings

From the 61 papers which studied similar peatland ecosystem properties as the EMBER project publications, the ecosystem properties with the largest number of relevant papers were those concerned with alterations to soil physical and chemical properties, and *Sphagnum* growth/abundance (Table 4). So far, there appears to be a consensus in the literature for four ecosystem properties and their response to burning: (1) mean and/or maximum soil temperatures increase following prescribed burning, (2) prescribed burning is associated with increased exposure of the peat surface and/or more erosion, (3) prescribed burning alters catchment hydrological functions, and (4) prescribed burning reduces aquatic invertebrate diversity. However, the latter only included studies undertaken by the EMBER research group. A majority of the available studies have also reported some form of alteration to soil physical and/or chemical properties, and stream chemistry following prescribed burning. Results suggest that there is the greatest level of variability in conclusions of studies that have examined burning effects on *Sphagnum* growth/abundance. The overall net effect (for the first four ecosystem properties) shows that, to date, >5-fold more studies have reported negative effects of moorland vegetation burning on the environment compared to positive effects, and with almost a 4-fold number of negative effects reported compared to no/mixed effects.

### 3.3 Potential sponsorship bias

Of the 61 papers summarised in Table 4, eight had declared funding links to grouse industry groups, while 24 were funded by government agencies that form national-scale environmental policy (Supplementary Table 5). While there were no clear links between government agency funding and any research outcomes, for studies funded by grouse industry groups there was a significantly higher probability of reporting positive effects of burning, most notably for *Sphagnum* growth/abundance response to burning (*p=0.00073)*, but also for the overall net effect conclusions (*p=0.00015*; Table 5, Supplementary Table 6). For *Sphagnum* growth/abundance specifically, four papers with grouse industry funding links have suggested positive effects but none have suggested any negative effects, in contrast to 6 negatives and 5 no/mixed effects funded by non-grouse shooting organisations. All four of those positive papers (Lee et al., 2013, Marrs et al., 2019b, Milligan et al., 2018, Harris et al., 2011) are from the same research group, whose long-time head is the President of the Heather Trust, a pro-burning, grouse industry group. Three of those four papers reported results from the same experiment undertaken at the Hard Hill burning plots at Moor House, northern England. Notably, the two most recent of these papers made prominent claims about heather burning being beneficial for mitigating wildfire risk, yet such effects were not studied in either paper.

**Table 5.** Fisher’s Test for grouse industry and government funded projects, versus those funded by other groups.

| **Funding/**<br>**Ecosystem property** | **Overall test $p$** | **positive:**<br>**no-effect** | **positive:**<br>**negative** | **No-effect:**<br>**negative** |
| --- | --- | --- | --- | --- |
| ***Grouse industry funding*** |  |  |  |  |
| *Sphagnum*<br>growth/abundance | 0.0007326 | 0.01429 | 0.02381 | 1 |
| Soil temperature | 1 | 1 | 1 | 1 |
| Bare peat/erosion | 1 | 1 | 1 | 1 |
| Aquatic invertebrates | 1 | 1 | 1 | 1 |
| Hydrological function | 0.4857 |  |  |  |
| Stream chemistry | 1 |  |  |  |
| Peat physical/<br>chemical properties | 1 |  |  |  |
| Net effect | 0.0001475 | 0.01429 | 0.0008547 | 1 |
| ***Government funding*** |  |  |  |  |
| *Sphagnum*<br>growth/abundance | 0.6743 | 1 | 1 | 1 |
| Soil temperature | 1 | 1 | 1 | 1 |
| Bare peat/erosion | 1 | 1 | 1 | 1 |
| Aquatic invertebrates | 1 | 1 | 1 | 1 |
| Hydrological function | 1 |  |  |  |
| Stream chemistry | 0.4 |  |  |  |
| Peat physical/<br>chemical properties | 1 |  |  |  |
| Net effect | 0.1903 | 1 | 0.3846 | 1 |

Sponsorship bias is a known phenomenon in scientific research (Lesser et al., 2007), although this is not necessarily problematic if researchers are open and transparent about their reasons for undertaking a piece of research. More problematic, given our results, could be a situation where researchers failed to publicise their funders or conflicts of interest, because this might have the potential to create a situation in which an organisation can remain hidden, yet gains support for their cause, behind a veneer of ‘independent’ scientific critique.

We have concerns about the potential for undeclared funding and other conflicts of interest that may entirely undermine the conclusions of some moorland burning research studies (e.g. Marrs et al., 2019a). In the version of their EMBER commentary paper that was subject to peer review and editorial decision, A&H declared no funding for their work. In the final version produced after acceptance they modified the statement to acknowledge Natural Environment Research Council funding, but some of this (UK PopNet, 2006-2010) appears to have ended several years before any EMBER paper was even published. Of greater concern, the second author, Heinemeyer, received a grant of £25,000 in 2017 from the British Association for Shooting and Conservation (BASC), a pro-burning, pro-grouse shooting, gun sports association. These funding links are clearly documented online (https://basc.org.uk/blog/uncategorized/basc-backs-moorland-study/, last accessed 29 July 2019) and were also reported in the Shooting Times (June 14^th^ 2017, p7). Funding from BASC or any other pro-grouse shooting organisation of course does not disqualify the authors from criticising other research studies. However, the BASC announcement of the funding to Heinemeyer also, somewhat strangely, included criticism of the EMBER project attributed to named BASC staff. It is not clear why A&H feel that these links should not be declared and explained so that readers can make a fully informed judgement of their critique.

A&H further cited in their conclusions that there is forthcoming work from Heinemeyer et al. (the document is not available) about a project called Peatland-ES-UK. They infer that this will provide new evidence, and hence that the EMBER project findings should be revisited by policy-makers. According to the Peatland-ES-UK project website http://peatland-es-uk.york.ac.uk, last accessed 29 July 2019) Phase 2 of that project is currently funded by the Moorland Association (another pro-burning, gun-sports lobby group) as well as BASC. On 15 October 2018, Heinemeyer also published, on social media, a note of thanks to the Heather Trust for funding the Peatland-ES-UK project (https://twitter.com/AndreasHeinem/status/1051819786265616384, last accessed 29 July 2019). The EMBER commentary paper (Ashby and Heinemeyer, 2019) was listed by Heinemeyer as an apparent output from Peatland-ES-UK at the project’s advisory group meeting in March 2019, suggesting that Heinemeyer considered it part of the project. Thus, it is unclear why A&H also failed to declare these funding sources within their paper’s acknowledgements at the time of submission.

## 4. Conclusion

In this paper we have shown that: (i) geographical variability did not confound EMBER results; (ii) the EMBER soil temperature findings are robust, and; (iii) a systematic review of the literature illustrates that the EMBER project findings are broadly in line with the majority of other published studies on similar studied variables impacted by prescribed moorland burning. We therefore dismiss the assertions by A&H. We have also shown that some UK studies that suggest moorland burning is beneficial for the environment have funding links to pro-grouse shooting, pro-burning organisations. Policy-makers need sound evidence to support the policy process on moorland burning. While funding sources and potential conflicts of interest should always be declared from the outset of the peer-review and publication process, unfortunately we have shown this is not always the case.

## Acknowledgements

Our research on peatland management as part of the EMBER project was funded by NERC (NE/G00224X/1) and Yorkshire Water. We have also received recent funding and in-kind support for our upland research more generally from numerous organisations with upland interests, including Calderdale Council, Environment Agency, Forestry Commission, JBA Trust, Defra, the Moorland Association, Moors for the Future/Peak District NPA, Natural England, North Pennines AONB/Durham County Council, RSPB, The National Trust, UK PopNet, United Utilities, Dwr Cymru Welsh Water, and the Yorkshire Dales Rivers Trust. The Game & Wildlife Conservation Trust and R.AG.T. Seeds are members of the advisory board for two NERC/BBSRC/Defra/Scottish Government-funded lowland farm/soil management projects (NE/M017079/1, BB/L026023/1) involving Holden. Brown currently works on a project supported in-kind by a grouse moor owner in the Yorkshire Dales to collect evidence regarding the efficacy of natural flood management techniques. Holden leads the NERC iCASP project (NE/P011160/1) which includes a range of additional partner organisations who provide in-kind support (https://icasp.org.uk/partners/).

## Author contributions

Both authors contributed equally to this piece of work.

## Data accessibility

Relevant data, or links to larger files, are provided in the supplementary material.

## Supplementary information

**Supplementary Table 1.** Mean water temperature data at the ten EMBER sites studied in Brown et al. (2013) collected between spring 2010 and autumn 2011.

| **River** | **Mean water temperature (°C)** |
| --- | --- |
| Bull Clough | 9.92 |
| Crowden Little Brook | 11.96 |
| Great Egglehope Beck | 8.85 |
| Green Burn | 8.92 |
| Lodgegill Syke | 8.19 |
| Moss Burn | 9.94 |
| Oakner Clough | 10.84 |
| Rising Clough | 11.82 |
| Trout Beck | 9.07 |
| Woo Gill | 10.11 |

**Supplementary Table 2.** Fine particulate organic matter density and monthly rainfall data (based on the UKCIP09 5 km gridded model data presented by A&H) for the ten EMBER rivers studied in Brown et al. (2013).

| River | Mean monthly rainfall<br>(mm) | FPOM (g m<sup>2</sup>) |
| --- | --- | --- |
| Bull Clough | 57.18 | 5.06 |
|  | 118.3 | 2.01 |
|  | 16.79 | 4.35 |
|  | 85.13 | 5.03 |
|  | 106.68 | 8.3 |
| Great Egglesthorpe Beck | 54.02 | 18.21 |
|  | 80.27 | 0.78 |
|  | 19.15 | 102.95 |
|  | 53.62 | 2.88 |
|  | 78.27 | 1.87 |
| Lodgegill Syke | 54.02 | 3.5 |
|  | 80.27 | 1.72 |
|  | 19.15 | 6.14 |
|  | 53.62 | 2.93 |
|  | 78.27 | 3.72 |
| Rising Clough | 92.64 | 5.49 |
|  | 94.24 | 37.92 |
|  | 15.03 | 5.28 |
|  | 71.15 | 6.1 |
|  | 65.02 | 4.92 |
| Woo Gill | 61.61 | 2.01 |
|  | 106.62 | 0.33 |
|  | 31.19 | 9.7 |
|  | 83.87 | 1.24 |
|  | 111.07 | 132.98 |
| Crowden Little Brook | 53.77 | 0.71 |
|  | 123.06 | 0.73 |
|  | 25.81 | 1.24 |
|  | 97.24 | 1.6 |
|  | 139.43 | 1.65 |
| Green Burn | 50.33 | 6.59 |
|  | 140.11 | 2.34 |
|  | 77.37 | 3.19 |
|  | 100.04 | 2.48 |
|  | 218.65 | 1.61 |
| Moss Burn | 50.33 | 2.04 |
|  | 140.11 | 2.18 |
|  | 77.37 | 4.24 |
|  | 100.04 | 4.16 |
|  | 218.65 | 2.96 |
| Oakner Clough | 47.36 | 1.84 |
|  | 115.55 | 0.58 |
|  | 26.5 | 1.1 |
| Trout Beck | 59.91 | 0.47 |
|  | 140.21 | 11.2 |
|  | 51.97 | 3.41 |
|  | 117.68 | 3.47 |
|  | 178.36 | 2.65 |

**Supplementary Table 3.** Daily maximum soil surface temperature disturbances. Top 10 % of values reported in Brown et al. (2015) have been removed. Numbers represent burn plot age. Low/med/high refers to the topographic index location of each plot

| 15low | 7low | 4low | 2low | 15med | 7med | 4med | 2med | 15high | 7high | 4high | 2high |
| --- | --- | --- | --- | --- | --- | --- | --- | --- | --- | --- | --- |
| 2.48 | 3.25 | 6.66 | 9.03 | 2.07 | 2.38 | 5.79 | 4.35 | 2.44 | 4.91 | 2.89 | 8.07 |
| 2.38 | 3.17 | 6.55 | 8.98 | 2.01 | 2.32 | 5.76 | 4.26 | 2.43 | 4.90 | 2.87 | 7.87 |
| 2.24 | 3.04 | 6.33 | 8.96 | 2.01 | 2.30 | 5.73 | 4.21 | 2.29 | 4.76 | 2.65 | 7.74 |
| 2.13 | 2.91 | 6.12 | 8.80 | 1.92 | 2.20 | 5.67 | 4.16 | 2.23 | 4.67 | 2.62 | 7.66 |
| 2.11 | 2.85 | 5.91 | 8.70 | 1.92 | 2.18 | 5.67 | 4.12 | 2.23 | 4.63 | 2.59 | 7.34 |
| 2.10 | 2.75 | 5.81 | 8.65 | 1.90 | 2.13 | 5.57 | 4.12 | 2.22 | 4.61 | 2.53 | 7.24 |
| 2.10 | 2.75 | 5.64 | 8.56 | 1.85 | 1.97 | 5.50 | 3.64 | 2.09 | 4.03 | 2.39 | 7.23 |
| 2.09 | 2.69 | 5.48 | 7.65 | 1.77 | 1.95 | 5.44 | 3.64 | 2.08 | 3.93 | 2.33 | 6.83 |
| 2.08 | 2.53 | 5.30 | 7.41 | 1.77 | 1.75 | 5.37 | 3.60 | 2.05 | 3.59 | 2.15 | 6.81 |
| 2.00 | 2.47 | 5.20 | 7.38 | 1.64 | 1.63 | 5.17 | 3.52 | 1.94 | 3.38 | 2.03 | 6.80 |
| 1.99 | 2.39 | 5.03 | 7.31 | 1.59 | 1.48 | 4.95 | 3.51 | 1.93 | 3.36 | 1.98 | 6.47 |
| 1.78 | 2.35 | 4.86 | 7.19 | 1.57 | 1.43 | 4.77 | 3.44 | 1.84 | 3.27 | 1.97 | 6.44 |
| 1.69 | 2.34 | 4.70 | 7.14 | 1.50 | 1.41 | 4.01 | 3.38 | 1.79 | 3.23 | 1.94 | 6.26 |
| 1.63 | 2.16 | 4.68 | 6.85 | 1.50 | 1.37 | 3.96 | 3.37 | 1.74 | 3.17 | 1.85 | 5.79 |
| 1.60 | 2.13 | 4.52 | 6.69 | 1.49 | 1.33 | 3.74 | 3.33 | 1.74 | 3.04 | 1.83 | 5.54 |
| 1.60 | 2.12 | 4.52 | 6.67 | 1.38 | 1.29 | 3.65 | 3.22 | 1.73 | 3.02 | 1.82 | 5.47 |
| 1.60 | 2.07 | 4.37 | 6.51 | 1.35 | 1.20 | 3.61 | 3.11 | 1.61 | 2.67 | 1.78 | 5.45 |
| 1.57 | 2.07 | 4.23 | 6.30 | 1.32 | 1.15 | 3.58 | 2.91 | 1.61 | 2.57 | 1.73 | 5.30 |
| 1.56 | 2.07 | 4.18 | 6.20 | 1.29 | 1.12 | 3.42 | 2.90 | 1.60 | 2.52 | 1.72 | 5.28 |
| 1.55 | 2.03 | 3.99 | 6.19 | 1.26 | 1.06 | 3.16 | 2.79 | 1.58 | 2.51 | 1.65 | 4.74 |
| 1.48 | 1.95 | 3.91 | 5.72 | 1.24 | 1.04 | 3.13 | 2.67 | 1.57 | 2.46 | 1.64 | 4.52 |
| 1.31 | 1.90 | 3.89 | 5.71 | 1.21 | 0.98 | 2.98 | 2.48 | 1.57 | 2.42 | 1.57 | 4.38 |
| 1.30 | 1.85 | 3.82 | 5.69 | 1.16 | 0.97 | 2.97 | 2.45 | 1.56 | 2.41 | 1.48 | 4.36 |
| 1.25 | 1.83 | 3.69 | 5.57 | 1.16 | 0.97 | 2.92 | 2.43 | 1.50 | 2.40 | 1.46 | 4.32 |
| 1.23 | 1.79 | 3.61 | 5.54 | 1.16 | 0.94 | 2.83 | 2.25 | 1.49 | 2.31 | 1.35 | 4.00 |
| 1.20 | 1.62 | 3.54 | 4.95 | 1.08 | 0.88 | 2.80 | 2.15 | 1.35 | 2.25 | 1.35 | 3.67 |
| 1.17 | 1.50 | 3.51 | 4.87 | 1.00 | 0.87 | 2.62 | 2.12 | 1.29 | 2.16 | 1.28 | 3.52 |
| 1.13 | 1.41 | 3.30 | 4.73 | 0.99 | 0.82 | 2.56 | 2.03 | 1.23 | 2.16 | 1.27 | 3.44 |
| 1.07 | 1.40 | 3.28 | 4.63 | 0.90 | 0.75 | 2.51 | 2.02 | 1.21 | 2.09 | 1.25 | 3.23 |
| 1.04 | 1.34 | 3.28 | 4.48 | 0.83 | 0.73 | 2.48 | 1.98 | 1.14 | 2.06 | 1.24 | 3.21 |
| 1.03 | 1.30 | 3.27 | 4.32 | 0.79 | 0.70 | 2.31 | 1.95 | 1.03 | 2.04 | 1.15 | 3.18 |
| 1.01 | 1.29 | 3.23 | 4.17 | 0.75 | 0.67 | 2.26 | 1.93 | 0.93 | 1.96 | 1.14 | 3.17 |
| 0.91 | 1.29 | 3.23 | 4.07 | 0.72 | 0.67 | 2.26 | 1.93 | 0.91 | 1.87 | 1.09 | 3.04 |
| 0.90 | 1.28 | 3.14 | 4.02 | 0.71 | 0.56 | 2.19 | 1.91 | 0.88 | 1.85 | 1.04 | 2.92 |
| 0.90 | 1.23 | 3.11 | 3.72 | 0.70 | 0.52 | 2.11 | 1.83 | 0.87 | 1.74 | 0.98 | 2.90 |
| 0.78 | 1.23 | 3.11 | 3.70 | 0.67 | 0.51 | 2.06 | 1.78 | 0.81 | 1.72 | 0.89 | 2.88 |
| 0.77 | 1.17 | 3.09 | 3.68 | 0.65 | 0.51 | 2.05 | 1.76 | 0.79 | 1.67 | 0.84 | 2.87 |
| 0.72 | 1.09 | 3.08 | 3.61 | 0.65 | 0.46 | 2.05 | 1.71 | 0.79 | 1.62 | 0.76 | 2.80 |
| 0.70 | 1.03 | 3.08 | 3.45 | 0.62 | 0.41 | 1.94 | 1.66 | 0.75 | 1.60 | 0.73 | 2.80 |
| 0.67 | 0.98 | 2.92 | 3.44 | 0.56 | 0.40 | 1.88 | 1.63 | 0.73 | 1.54 | 0.68 | 2.76 |
| 0.64 | 0.98 | 2.88 | 3.43 | 0.53 | 0.34 | 1.83 | 1.62 | 0.73 | 1.51 | 0.62 | 2.66 |
| 0.64 | 0.97 | 2.71 | 3.43 | 0.43 | 0.32 | 1.81 | 1.60 | 0.72 | 1.36 | 0.49 | 2.63 |
| 0.58 | 0.92 | 2.68 | 3.39 | 0.42 | 0.27 | 1.81 | 1.54 | 0.71 | 1.27 | 0.47 | 2.60 |
| 0.58 | 0.90 | 2.67 | 3.27 | 0.40 | 0.27 | 1.78 | 1.53 | 0.67 | 1.25 | 0.47 | 2.53 |
| 0.56 | 0.88 | 2.57 | 3.17 | 0.34 | 0.25 | 1.73 | 1.51 | 0.63 | 1.23 | 0.43 | 2.50 |
| 0.53 | 0.83 | 2.56 | 3.12 | 0.33 | 0.23 | 1.73 | 1.49 | 0.57 | 1.16 | 0.41 | 2.31 |
| 0.46 | 0.83 | 2.56 | 3.09 | 0.32 | 0.21 | 1.63 | 1.35 | 0.57 | 1.07 | 0.40 | 2.29 |
| 0.41 | 0.80 | 2.56 | 3.08 | 0.30 | 0.20 | 1.62 | 1.34 | 0.54 | 0.84 | 0.29 | 2.19 |
| 0.35 | 0.79 | 2.50 | 3.05 | 0.29 | 0.20 | 1.62 | 1.30 | 0.53 | 0.83 | 0.26 | 2.14 |
| 0.35 | 0.76 | 2.44 | 2.80 | 0.27 | 0.18 | 1.60 | 1.25 | 0.44 | 0.81 | 0.23 | 2.12 |
| 0.33 | 0.74 | 2.43 | 2.75 | 0.26 | 0.18 | 1.58 | 1.25 | 0.37 | 0.79 | 0.23 | 2.10 |
| 0.32 | 0.67 | 2.31 | 2.64 | 0.25 | 0.18 | 1.56 | 1.24 | 0.35 | 0.72 | 0.19 | 2.07 |
| 0.29 | 0.66 | 2.17 | 2.55 | 0.22 | 0.18 | 1.52 | 1.24 | 0.34 | 0.67 | 0.18 | 2.06 |
| 0.26 | 0.66 | 2.16 | 2.54 | 0.22 | 0.17 | 1.49 | 1.21 | 0.34 | 0.61 | 0.16 | 2.05 |
| 0.23 | 0.62 | 2.05 | 2.46 | 0.21 | 0.15 | 1.43 | 1.19 | 0.24 | 0.60 | 0.14 | 2.04 |
| 0.22 | 0.62 | 2.04 | 2.43 | 0.21 | 0.14 | 1.37 | 1.17 | 0.24 | 0.51 | 0.13 | 2.02 |
| 0.21 | 0.59 | 1.97 | 2.17 | 0.20 | 0.12 | 1.35 | 1.08 | 0.22 | 0.49 | 0.13 | 1.77 |
| 0.14 | 0.59 | 1.74 | 2.08 | 0.20 | 0.12 | 1.28 | 1.07 | 0.20 | 0.42 | 0.11 | 1.74 |
| 0.00 | 0.57 | 1.73 | 1.98 | 0.19 | 0.09 | 1.24 | 1.05 | 0.17 | 0.42 | 0.09 | 1.64 |
| -0.05 | 0.51 | 1.70 | 1.97 | 0.18 | 0.08 | 1.23 | 1.03 | 0.16 | 0.40 | 0.06 | 1.63 |
| -0.07 | 0.47 | 1.63 | 1.96 | 0.16 | 0.06 | 1.22 | 0.94 | 0.15 | 0.38 | 0.05 | 1.63 |
| -0.07 | 0.46 | 1.63 | 1.93 | 0.16 | 0.06 | 1.13 | 0.91 | 0.10 | 0.34 | 0.05 | 1.61 |
| -0.09 | 0.46 | 1.61 | 1.89 | 0.13 | 0.05 | 1.08 | 0.90 | 0.08 | 0.32 | 0.01 | 1.56 |
| -0.11 | 0.44 | 1.56 | 1.82 | 0.12 | 0.02 | 0.99 | 0.83 | 0.04 | 0.32 | 0.01 | 1.55 |
| -0.12 | 0.40 | 1.56 | 1.77 | 0.11 | 0.02 | 0.94 | 0.82 | 0.02 | 0.32 | -0.05 | 1.49 |
| -0.12 | 0.39 | 1.54 | 1.74 | 0.10 | -0.01 | 0.92 | 0.74 | 0.01 | 0.28 | -0.09 | 1.48 |
| -0.12 | 0.34 | 1.49 | 1.69 | 0.10 | -0.06 | 0.85 | 0.73 | 0.00 | 0.26 | -0.10 | 1.41 |
| -0.16 | 0.33 | 1.42 | 1.63 | 0.10 | -0.06 | 0.80 | 0.72 | 0.00 | 0.25 | -0.12 | 1.19 |
| -0.16 | 0.28 | 1.41 | 1.61 | 0.09 | -0.06 | 0.77 | 0.69 | -0.03 | 0.25 | -0.12 | 1.11 |
| -0.16 | 0.25 | 1.26 | 1.58 | 0.00 | -0.10 | 0.71 | 0.61 | -0.03 | 0.22 | -0.13 | 1.09 |
| -0.18 | 0.25 | 1.24 | 1.55 | 0.00 | -0.19 | 0.70 | 0.60 | -0.10 | 0.21 | -0.14 | 0.99 |
| -0.20 | 0.22 | 1.23 | 1.53 | -0.01 | -0.20 | 0.66 | 0.58 | -0.10 | 0.19 | -0.17 | 0.94 |
| -0.23 | 0.20 | 1.23 | 1.49 | -0.02 | -0.21 | 0.57 | 0.48 | -0.10 | 0.10 | -0.20 | 0.88 |
| -0.24 | 0.17 | 1.21 | 1.40 | -0.02 | -0.21 | 0.56 | 0.46 | -0.14 | 0.05 | -0.20 | 0.86 |
| -0.28 | 0.16 | 1.14 | 1.35 | -0.03 | -0.21 | 0.56 | 0.44 | -0.14 | 0.04 | -0.20 | 0.81 |
| -0.30 | 0.13 | 1.12 | 1.30 | -0.06 | -0.23 | 0.54 | 0.36 | -0.15 | 0.03 | -0.21 | 0.81 |
| -0.31 | 0.11 | 1.08 | 1.26 | -0.11 | -0.34 | 0.39 | 0.35 | -0.15 | 0.01 | -0.22 | 0.81 |
| -0.32 | 0.09 | 1.07 | 1.23 | -0.13 | -0.38 | 0.34 | 0.33 | -0.16 | -0.01 | -0.22 | 0.74 |
| -0.33 | 0.06 | 1.02 | 1.18 | -0.14 | -0.40 | 0.34 | 0.33 | -0.18 | -0.03 | -0.23 | 0.65 |
| -0.36 | 0.00 | 1.02 | 1.16 | -0.19 | -0.41 | 0.29 | 0.32 | -0.22 | -0.03 | -0.23 | 0.64 |
| -0.37 | -0.01 | 1.01 | 1.05 | -0.21 | -0.42 | 0.28 | 0.28 | -0.24 | -0.07 | -0.25 | 0.51 |
| -0.39 | -0.02 | 0.96 | 1.01 | -0.21 | -0.45 | 0.22 | 0.28 | -0.24 | -0.12 | -0.25 | 0.47 |
| -0.45 | -0.02 | 0.87 | 1.01 | -0.32 | -0.46 | 0.20 | 0.27 | -0.25 | -0.16 | -0.28 | 0.44 |
| -0.46 | -0.07 | 0.76 | 1.00 | -0.32 | -0.47 | 0.20 | 0.25 | -0.26 | -0.18 | -0.29 | 0.43 |
| -0.48 | -0.08 | 0.73 | 0.98 | -0.35 | -0.48 | 0.19 | 0.24 | -0.28 | -0.24 | -0.29 | 0.40 |
| -0.50 | -0.09 | 0.71 | 0.92 | -0.35 | -0.49 | 0.16 | 0.23 | -0.30 | -0.26 | -0.31 | 0.40 |
| -0.52 | -0.09 | 0.69 | 0.86 | -0.36 | -0.51 | 0.16 | 0.21 | -0.30 | -0.28 | -0.35 | 0.29 |
| -0.53 | -0.12 | 0.67 | 0.82 | -0.38 | -0.51 | 0.13 | 0.19 | -0.32 | -0.30 | -0.38 | 0.27 |
| -0.54 | -0.14 | 0.49 | 0.80 | -0.39 | -0.52 | 0.13 | 0.18 | -0.34 | -0.30 | -0.39 | 0.24 |
| -0.56 | -0.15 | 0.36 | 0.73 | -0.41 | -0.54 | 0.13 | 0.15 | -0.37 | -0.30 | -0.40 | 0.23 |
| -0.58 | -0.16 | 0.35 | 0.72 | -0.42 | -0.56 | 0.05 | 0.13 | -0.38 | -0.33 | -0.44 | 0.22 |
| -0.66 | -0.27 | 0.34 | 0.63 | -0.43 | -0.58 | 0.05 | 0.10 | -0.41 | -0.34 | -0.45 | 0.18 |
| -0.66 | -0.30 | 0.23 | 0.62 | -0.44 | -0.63 | 0.01 | 0.08 | -0.45 | -0.34 | -0.46 | 0.12 |
| -0.72 | -0.35 | 0.20 | 0.60 | -0.45 | -0.67 | -0.01 | 0.05 | -0.51 | -0.35 | -0.49 | 0.07 |
| -0.74 | -0.37 | 0.20 | 0.52 | -0.51 | -0.69 | -0.09 | 0.05 | -0.55 | -0.38 | -0.54 | -0.02 |
| -0.81 | -0.41 | 0.16 | 0.38 | -0.54 | -0.71 | -0.10 | 0.04 | -0.58 | -0.44 | -0.56 | -0.04 |
| -0.84 | -0.41 | -0.12 | 0.37 | -0.54 | -0.71 | -0.14 | 0.04 | -0.64 | -0.46 | -0.60 | -0.10 |
| -0.85 | -0.41 | -0.12 | 0.35 | -0.61 | -0.73 | -0.17 | 0.01 | -0.64 | -0.47 | -0.60 | -0.13 |
| -0.88 | -0.48 | -0.15 | 0.26 | -0.62 | -0.74 | -0.19 | 0.00 | -0.71 | -0.51 | -0.62 | -0.20 |
| -0.88 | -0.51 | -0.17 | 0.25 | -0.67 | -0.74 | -0.23 | -0.03 | -0.74 | -0.52 | -0.62 | -0.22 |
| -0.91 | -0.54 | -0.18 | 0.22 | -0.68 | -0.79 | -0.28 | -0.08 | -0.74 | -0.55 | -0.62 | -0.24 |
| -0.91 | -0.58 | -0.20 | 0.21 | -0.69 | -0.79 | -0.30 | -0.09 | -0.75 | -0.56 | -0.62 | -0.26 |
| -0.92 | -0.60 | -0.20 | 0.17 | -0.71 | -0.83 | -0.34 | -0.12 | -0.82 | -0.60 | -0.63 | -0.26 |
| -0.94 | -0.62 | -0.23 | 0.05 | -0.76 | -0.85 | -0.36 | -0.20 | -0.84 | -0.65 | -0.64 | -0.27 |
| -0.94 | -0.66 | -0.25 | 0.04 | -0.76 | -0.87 | -0.40 | -0.24 | -0.90 | -0.69 | -0.76 | -0.28 |
| -1.02 | -0.66 | -0.28 | 0.04 | -0.76 | -0.89 | -0.47 | -0.34 | -0.90 | -0.70 | -0.76 | -0.33 |
| -1.08 | -0.66 | -0.30 | -0.02 | -0.77 | -0.90 | -0.48 | -0.37 | -1.00 | -0.72 | -0.81 | -0.38 |
| -1.12 | -0.68 | -0.39 | -0.10 | -0.86 | -0.93 | -0.50 | -0.43 | -1.07 | -0.73 | -0.85 | -0.44 |
| -1.12 | -0.68 | -0.41 | -0.11 | -0.91 | -0.96 | -0.58 | -0.44 | -1.15 | -0.78 | -0.86 | -0.44 |
| -1.17 | -0.69 | -0.46 | -0.21 | -0.98 | -0.97 | -0.63 | -0.47 | -1.17 | -0.82 | -0.90 | -0.48 |
| -1.17 | -0.74 | -0.47 | -0.22 | -0.98 | -1.00 | -0.67 | -0.48 | -1.19 | -0.84 | -0.94 | -0.51 |
| -1.20 | -0.78 | -0.48 | -0.26 | -1.10 | -1.13 | -0.68 | -0.52 | -1.21 | -0.86 | -0.94 | -0.51 |
| -1.23 | -0.78 | -0.57 | -0.30 | -1.24 | -1.15 | -0.72 | -0.54 | -1.24 | -0.92 | -1.14 | -0.54 |
| -1.24 | -0.85 | -0.76 | -0.30 | -1.28 | -1.17 | -0.80 | -0.55 | -1.24 | -1.02 | -1.20 | -0.60 |
| -1.27 | -0.85 | -0.79 | -0.38 | -1.32 | -1.52 | -0.81 | -0.56 | -1.25 | -1.08 | -1.31 | -0.63 |
| -1.28 | -0.91 | -0.81 | -0.41 | -1.33 | -1.55 | -0.82 | -0.64 | -1.25 | -1.21 | -1.32 | -0.63 |
| -1.37 | -0.94 | -0.86 | -0.46 | -1.36 | -1.57 | -0.83 | -0.70 | -1.27 | -1.32 | -1.35 | -0.72 |
| -1.38 | -0.96 | -0.92 | -0.51 | -1.36 | -1.57 | -0.86 | -0.71 | -1.31 | -1.34 | -1.49 | -0.72 |
| -1.44 | -1.03 | -0.92 | -0.53 | -1.42 | -1.61 | -0.93 | -0.77 | -1.32 | -1.54 | -1.49 | -0.75 |
| -1.54 | -1.03 | -0.99 | -0.55 | -1.44 | -1.65 | -1.04 | -0.80 | -1.33 | -1.55 | -1.52 | -0.87 |
| -1.58 | -1.04 | -0.99 | -0.56 | -1.47 | -1.76 | -1.05 | -0.85 | -1.36 | -1.62 | -1.56 | -1.05 |
| -1.65 | -1.08 | -1.05 | -0.57 | -1.55 | -1.99 | -1.08 | -0.91 | -1.37 | -1.75 | -1.61 | -1.25 |
| -1.67 | -1.11 | -1.27 | -0.76 | -1.59 | -2.03 | -1.12 | -0.96 | -1.51 | -1.82 | -1.67 | -1.29 |
| -1.71 | -1.14 | -1.29 | -0.78 | -1.60 | -2.11 | -1.19 | -1.01 | -1.52 | -1.83 | -1.73 | -1.38 |
| -1.80 | -1.16 | -1.51 | -0.84 | -1.64 | -2.16 | -1.22 | -1.04 | -1.59 | -1.93 | -1.78 | -1.39 |
| -1.83 | -1.20 | -1.74 | -1.34 | -1.71 | -2.38 | -1.39 | -1.13 | -2.08 | -2.07 | -1.97 | -1.42 |
| -2.11 | -1.27 | -1.84 | -1.44 | -1.72 | -2.48 | -1.42 | -1.14 | -2.18 | -2.17 | -1.99 | -1.45 |
| -2.31 | -1.28 | -1.86 | -1.93 | -1.94 | -2.50 | -1.44 | -1.16 | -2.19 | -2.45 | -2.08 | -1.48 |
| -2.36 | -1.29 | -1.86 | -1.98 | -2.03 | -2.50 | -1.58 | -1.22 | -2.29 | -2.48 | -2.29 | -1.60 |
| -2.51 | -1.31 | -1.88 | -2.01 | -2.04 | -2.51 | -1.81 | -1.25 | -2.92 | -2.56 | -2.35 | -1.61 |
| -2.59 | -1.31 | -1.91 | -2.06 | -2.12 | -2.58 | -1.86 | -1.28 | -3.00 | -2.66 | -2.75 | -1.65 |
| -2.62 | -1.38 | -1.97 | -2.09 | -2.20 | -2.81 | -1.90 | -1.52 | -3.02 | -2.68 | -2.82 | -1.72 |
| -2.64 | -1.47 | -2.12 | -2.20 | -2.77 | -2.89 | -1.93 | -1.73 | -3.06 | -2.97 | -2.87 | -1.77 |
| -2.74 | -1.59 | -2.25 | -2.34 | -3.11 | -3.14 | -2.14 | -1.90 | -3.07 | -3.12 | -2.87 | -1.94 |
| -2.80 | -1.79 | -2.26 | -2.39 | -3.12 | -3.53 | -2.53 | -1.93 | -3.12 | -3.43 | -2.89 | -2.42 |
| -2.84 | -2.15 | -2.55 | -2.53 | -3.31 | -3.55 | -2.56 | -1.99 | -3.16 | -3.65 | -3.25 | -3.01 |
| -2.85 | -2.57 | -2.76 | -2.57 | -3.39 | -3.57 | -3.55 | -2.00 | -3.20 | -3.71 | -3.27 | -3.07 |
| -3.05 | -2.95 | -2.93 | -2.80 | -3.60 | -3.64 | -3.93 | -2.25 | -3.25 | -4.38 | -3.75 | -3.35 |
| -3.11 | -2.97 | -3.19 | -3.25 | -3.72 | -3.81 | -4.14 | -2.41 | -3.29 | -4.45 | -3.77 | -3.82 |
| -3.18 | -3.35 | -3.47 | -3.34 | -3.86 | -4.16 | -4.29 | -3.66 | -4.36 | -4.50 | -4.17 | -3.95 |
| -3.19 | -3.55 | -4.60 | -4.79 | -4.02 | -4.17 | -4.39 | -3.74 | -4.58 | -4.69 | -4.50 | -4.40 |
| -3.56 | -3.55 | -4.64 | -4.82 | -4.17 | -4.47 | -4.56 | -4.05 | -4.63 | -4.81 | -4.61 | -4.88 |
| -4.02 | -3.81 | -4.94 | -4.91 | -4.46 | -4.77 | -4.94 | -4.20 | -5.16 | -4.83 | -4.66 | -5.90 |
| -4.15 | -3.82 | -4.98 | -4.92 | -4.53 | -5.50 | -5.07 | -4.55 | -5.19 | -5.51 | -4.69 | -6.00 |
| -4.56 | -4.19 | -5.52 | -5.67 | -5.23 | -5.68 | -5.69 | -4.78 | -5.21 | -5.54 | -5.30 | -7.31 |
| -4.93 | -4.89 | -5.96 | -6.01 | -5.43 | -6.45 | -6.52 | -4.86 | -5.33 | -6.01 | -6.35 | -8.01 |
| -5.61 | -5.35 | -7.68 | -7.73 | -5.45 | -6.56 | -7.47 | -5.51 | -5.44 | -8.60 | -6.53 | -8.13 |

**Supplementary Table 4. List of papers considered as part of the systematic review** (available at https://doi.org/10.5518/623)

**Supplementary Table 5. Shortlist of papers considered to be relevant for comparison with the EMBER study**, including classifications for each ecosystem property, funding information, and funder group classifications (available at https://doi.org/10.5518/623)

**Supplementary Table 6.** Fisher’s exact test data, showing the number of studies per classification group against funding groups

|  |  |  |  |  |  |  |  |
| --- | --- | --- | --- | --- | --- | --- | --- |
| ***Sphagnum growth/abundance*** |  |  |  | ***Bare peat/erosion*** |  |  |  |
|  | **+** | **0** | **-** |  | **+** | **0** | **-** |
| *Grouse funded* | 4 | 0 | 0 | *Grouse funded* | 0 | 0 | 1 |
| *Not grouse funded* | 0 | 6 | 5 | *Not grouse funded* | 0 | 0 | 7 |
|  | **+** | **0** | **-** |  | **+** | **0** | **-** |
| *Policy groups funding* | 3 | 3 | 2 | *Policy groups funding* | 0 | 0 | 2 |
| *Other* | 1 | 3 | 3 | *Other* | 0 | 0 | 6 |
| ***Soil temperature*** |  |  |  | ***Aquatic invertebrates*** |  |  |  |
|  | **+** | **0** | **-** |  | **+** | **0** | **-** |
| *Grouse funded* | 0 | 0 | 0 | *Grouse funded* | 0 | 0 | 0 |
| *Not grouse funded* | 0 | 0 | 6 | *Not grouse funded* | 0 | 0 | 5 |
|  | **+** | **0** | **-** |  | **+** | **0** | **-** |
| *Policy groups funding* | 0 | 0 | 1 | *Policy groups funding* | 0 | 0 | 3 |
| *Other* | 0 | 0 | 5 | *Other* | 0 | 0 | 2 |
| ***Hydrological processes*** |  |  |  | ***Peat properties*** |  |  |  |
|  | **Y** | **N** |  |  | **Y** | **N** |  |
| *Grouse funded* | 0 | 0 |  | *Grouse funded* | 2 | 0 |  |
| *Not grouse funded* | 9 | 0 |  | *Not grouse funded* | 16 | 3 |  |
|  | **Y** | **N** |  |  | **Y** | **N** |  |
| *Policy groups funding* | 2 | 0 |  | *Policy groups funding* | 7 | 1 |  |
| *Other* | 7 | 0 |  | *Other* | 11 | 2 |  |
| ***Stream chemistry*** |  |  |  | ***Net effect*** |  |  |  |
|  | **Y** | **N** |  |  | **+** | **0** | **-** |
| *Grouse funded* | 0 | 0 |  | ***Grouse funded*** | 4 | 0 | 1 |
| *Not grouse funded* | 9 | 1 |  | ***Not grouse funded*** | 0 | 6 | 22 |
|  | **Y** | **N** |  |  | **+** | **0** | **-** |
| *Policy groups funding* | 3 | 1 |  | ***Policy groups funding*** | 3 | 3 | 7 |
| *Other* | 6 | 0 |  | ***Other*** | 1 | 3 | 16 |

**Supplementary Figure 1.**
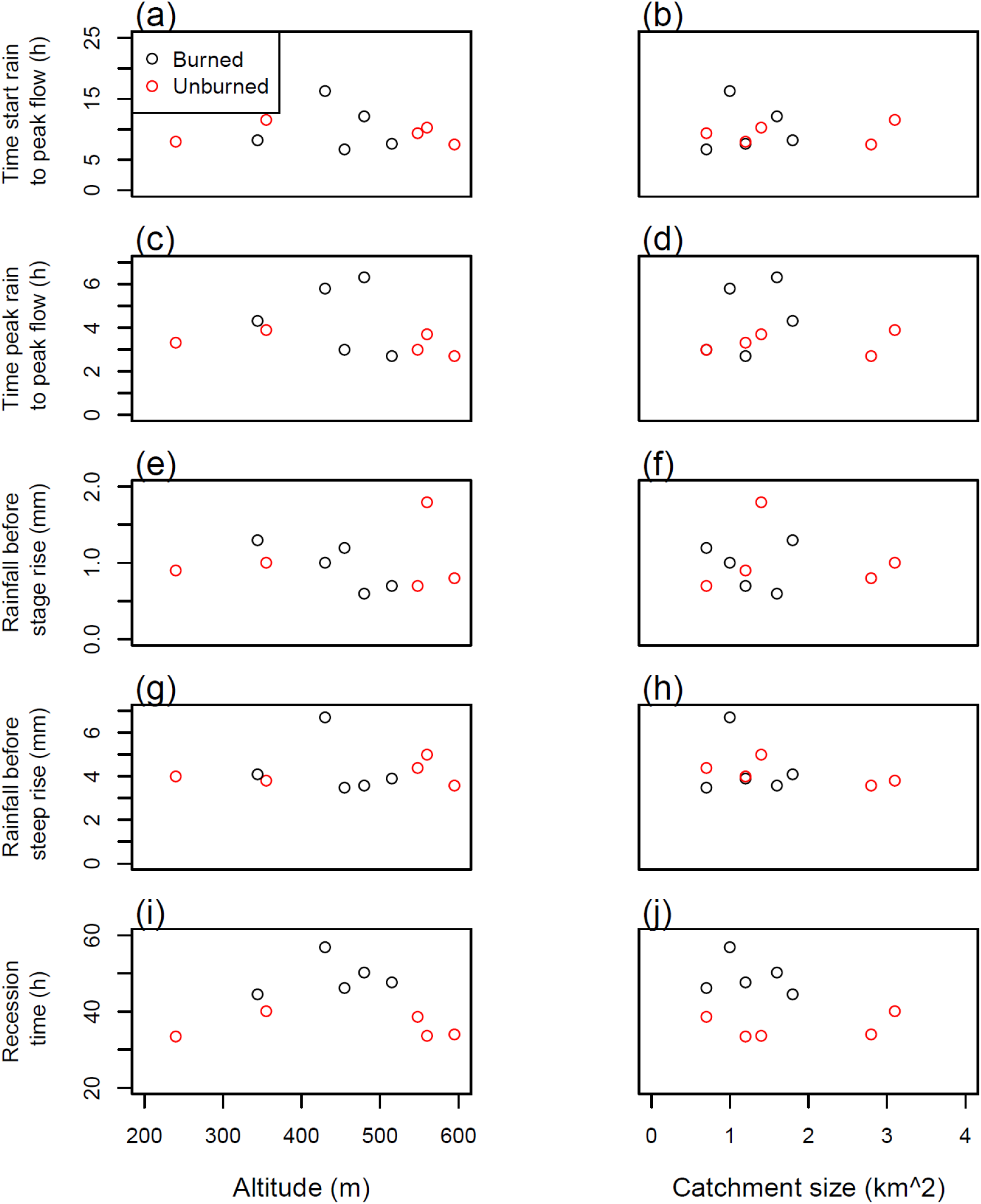
Scatterplots of hydrological metrics detailed in Holden et al. (2016) show no clear relationships with altitude or catchment size.

